# HIGH-RESOLUTION CHARACTERIZATION OF MALE ORNAMENTATION AND REEVALUATION OF SEX LINKAGE IN GUPPIES

**DOI:** 10.1101/2020.07.14.200329

**Authors:** Jake Morris, Iulia Darolti, Wouter van der Bijl, Judith E. Mank

## Abstract

Colouration plays a key role in the ecology of many species, influencing how an organism interacts with its environment, other species and conspecifics. Guppies are sexually dimorphic, with males displaying sexually selected colouration resulting from female preference. Previous work has suggested that much of guppy colour pattern variation is Y-linked. However, it remains unclear how many individual colour patterns are Y-linked in natural populations as much of the previous work has focused on phenotypes either not found in the wild, or aggregate measures such as total colour area. Moreover, ornaments have traditionally been identified and delineated by hand, and computational methods now make it possible to extract pixels and identify ornaments more automatedly, reducing the potential for human bias. Here we developed a pipeline for automated ornament identification and high-resolution image analysis of male guppy colour patterns and applied it to a multigenerational pedigree. Our results show that loci controlling the presence or absence of individual male ornaments in our population are not predominantly Y-linked. However, we find that ornaments of similar colour are not independent of each other, and modifier loci that affect whole animal colouration appear to be at least partially Y-linked. Considering these results, Y-linkage of individual ornaments may not be important in driving colour changes in natural populations of guppies, or in expansions of the non-recombining Y region, while Y-linked modifier loci that affect aggregate traits may well play an important role.

## Introduction

Colouration is an important morphological character in many species. It has the potential to greatly influence how an organism interacts with its environment via thermoregulation, UV protection [1–3], with other species via aposematism, mimicry and crypsis [4–7], and with conspecifics via intra-and intersexual selection [8,9]. Moreover, some colour phenotypes under sexual selection can incur significant metabolic production costs [10,11] or increased predation risks due to increased conspicuousness [12,13]. In these cases, the costs affect both sexes, but only one sex gains a mating advantage, resulting in sexual conflict [14,15]. In these cases, colour traits are likely to be dimorphic when the genetic architecture can be separated between females and males [15].

For species with male heterogametic sex chromosomes, one important way to establish separate male and female genetic architecture, and thereby resolve sexual conflict, is Y-linkage. Moreover, sexual conflict is thought to accelerate the evolution of the Y chromosome and its divergence from the X [16] when a sexually antagonistic allele is in close linkage with the sex-determining (Y) locus. In males, this will lead selection against recombinants to maintain linkage, ultimately resulting in a non-recombining Y chromosome region [17–19]. Theoretical models [16,17] predict an accumulation of sexually antagonistic genes on Y chromosomes, particularly in nascent sex chromosome systems before large-scale gene loss has occurred.

Male guppies (*Poecilia reticulata*) are brightly coloured and highly variable in their patterning [20]. These patterns are composed of spots and bars that can be categorized into three types, orange and red carotenoid and pterin ornaments, melanic black ornaments, and green and blue structural colours [21,22]. In contrast, females lack any of these ornaments and are instead a uniform colour. Although colouration leads to increased predation [23], male guppy colouration is the result of sexual selection, with females showing a preference for more colourful, brightly patterned males [20,21,24–26]. Guppies face fewer predators in the upper reaches of the streams and in these more recently colonised upstream populations, guppies have evolved more conspicuous colouration due to increased female preference [27,28]. Male guppy colouration is therefore a sexually antagonistic trait, being selectively disadvantageous via predation if it were found in females, but being sexually advantageous for males.

Previous work has suggested that a sizable proportion of guppy male colouration is Y-linked, and a summary by Lindholm and Breden (2002) identified 16 male ornament characters as fully Y-linked, 24 that recombine between the X and Y and two that are X linked. Indeed, Y-linkage of male colour traits in this system has been suggested as a main factor in the evolution of recombination suppression [29] between both the small region of the sex chromosomes that never recombines, as well as a larger region of the sex chromosomes where recombination occurs infrequently [30]. However, as Lindholm and Breden (2002) point out, much of the evidence for this Y-linkage has come from studies on phenotypes not typically found in the wild [31,32], or from studies that do not or cannot account for potential for variation in female sensitivity to hormone treatments [32–34], and/or from studies using aggregate measures of total colour area rather than specific ornaments [21,34,35]. Moreover, ornaments have traditionally been identified and delineated by hand, leading to the potential for human bias if samples are not analysed blind to parentage or place of origin [36].

Here we developed a pipeline for high resolution image analysis of male guppy colour patterns. We use this pipeline along with two datasets from our outbred stock population derived from the Quare river system to; i) identify and characterize male colour ornaments agnostically with computational clustering methods, ii) determine how these ornaments relate to each other, and iii) use pedigreed families to compare and contrast inheritance patterns and possible Y-linkage of both individual ornament phenotypes and aggregate pattern phenotypes of total colour area.

## Methods

Our stock population of guppies originates from the Quare River in Trinidad, originally collected in 1998 and subsequently maintained in large numbers (>500) in benign freely-mating conditions [37]. We established pedigrees from this population in 2016 by pairing four males and virgin females randomly in the first generation. We then paired offspring in subsequent generations from different families to maximise the number of grandparents in each brood, to minimize inbreeding which can make it difficult to differentiate different modes of inheritance, and to help increase the diversity of different phenotypes segregating in different families. This led to a minimum of six great-grandparents per family in generation three (see Supplementary Fig. 1). In all, our pedigree stretched across three generations and 19 crosses, and produced 214 male offspring across generations two and three (average per family = 14). We allowed between 4 and 8 months for each generation to permit multiple clutches in most pairs. We maintained both the stock population and pedigree families on a 12:12 light cycle at 26C and fed a standard hobby guppy diet and artemia daily.

### Imaging

We photographed males from the controlled pedigrees, described above, as well as 171 males taken from the main stock population and not used in pedigrees as a population baseline for ornamentation. For all males, fish were photographed as soon as possible after sexual maturation in a specially designed tank (photarium) with internal dimensions of Length 100mm x Width 100mm x Depth 20mm. Photos were taken with a tripod-mounted Canon EOS 100D and an 18-55m lens, using ISO 400, shutter speed 1/640 and F4.5. Lighting was standardized using a Polaroid PLLED312 312 LED light set to 5600K on full power. The standardized photographic conditions limited the need for pre-processing normalisation, however, we used the R function histMatch() from the RStoolbox v0.2.3 package [38] to correct the small amount of variation we did observe in brightness. In addition, we also used the R function focal() from the Raster v2.6-7 package [39] to slightly blur images for downstream colour block extraction, thereby eliminating any potential problems with glare or reflectance.

Following pre-processing, we cropped each photo to a 1801×931 (XY) pixel image using custom functions. We then employed a *k*-means clustering algorithm with function kmeans() from the package Stats v3.5.0 from the R core team [40], which partitions the image based on the RGB values of pixels into *k* clusters, so that the sum of squares from pixels to the assigned cluster RGB centres is minimized. Each image was partitioned using both *k*=20 and *k*=40, so that the image was split into 20 or 40 clusters of different pixels to highlight patches of same-coloured pixels that could be considered ornaments. *k*=20 and *k*=40 were chosen as these values provided a level of granularity that enabled reasonable extraction of ornaments across all fish in our dataset. With images now split into *k* clusters of different colours, we were then able to choose clusters that allowed us to separately extract orange and black pixels. For both orange and black pixels, we used *k*-means images to reduce the complexity of the colour on the fish to ease pixel cluster selection for the user. This was done from the *k*-image that gave the most distinct and defined pixel patterns. We did however extract the original RGB values from the non *k*-means image.

Following the extraction of orange and black pixels for each fish, we used geometric morphometric methods to align both main population and pedigree fish against a reference shape. This method has previously been used in colour analysis R packages such as Patternize v0.0.1 [41]. We placed 99 semi-landmarks (equally spaced points along a curve) around the body outlines, ignoring caudal, dorsal and other fins, in the original photos using the TPS suite (tpsDig 2.17 and tpsUtil 1.56) [42,43]. The gpagen() function from the Geomorph v3.0.7 package [44] was used to calculate the mean shape of the 171 male fish from the main population, and we used this as the reference shape against which the images of extracted pixels from all pedigree males were aligned. This alignment was done using Thin Plate Spline warping as implemented by the computeTransform() and applyTransform() functions from the package Morpho v2.6 [45]. See Supplementary Fig. 1 for pipeline overview.

### Defining ornaments and ornament inheritance

After aligning the orange and black extracted images of all 171 males from the stock population, we had RGB scores for each pixel in each ornament in each fish. We then used these data to define and delimit the male ornaments in our stock population. Pixels were marked as part of an ornament based on the proportion of presence across fish. For a pixel to be marked as ornamental it had to be found in > 11% of all assayed fish for orange pixels and > 10% of all assayed fish for black pixels. This slightly higher threshold for orange pigmentation was needed in order to separate orange ornaments OII and OIII which were otherwise joined by a few black pixels (Fig 1a; 1b). We then used the function hclust() from the package Stats v3.5.0 to cluster pixels into a predefined number of distinct groups, defined as ornaments. For orange ornaments we used a predefined number of four ornaments, as this was very clearly the number of distinct orange regions found in fish. For black ornaments we tested clustering using 6,7 and 8 ornaments, with one of the black regions being the eye. From these results, it was clear that there were five main patches of black pigmentation, plus the eye and one very small region that we did not consider in downstream analyses. From this we were able to measure the minimum and maximum XY coordinates or four orange and five black ornaments, and determine the size, hue and saturation of each ornament in each individual.

**Fig 1.**
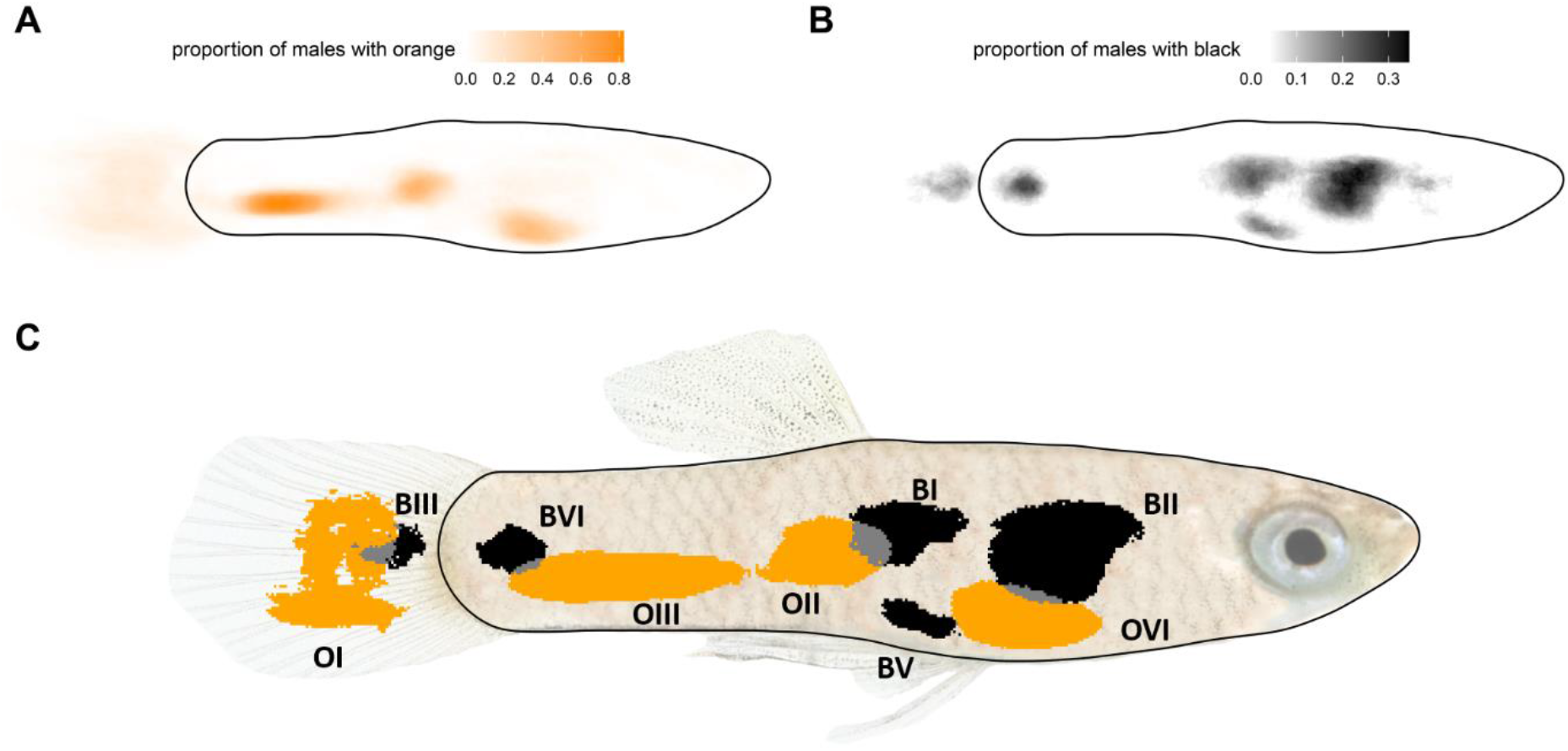
Male ornamentation. A) Heatmap showing the proportion of individuals from the full stock population with orange pigmentation at each pixel, B) Heatmap showing the proportion of individuals from the full stock population with black pigmentation at each pixel, with the eye masked, C) Mosaic plot showing the four orange and five black ornaments defined from the stock population. Grey pixels represent overlap between orange and black ornaments.

To better understand the correlational relationships among ornaments, referred to as modulation, we calculated Pearson’s correlations between ornaments for size (where ornaments with 10 pixels or less were marked as NA) and saturation in R using the rcorr.adjust() function (with use= “pairwise.complete.obs”) from the package RcmdrMisc v2.7-1 [46] which uses Holm’s method to apply a correction to P-values calculated using the rcorr() function from Hmisc [47]. We also calculated Spearman’s correlations for presence/absence (with ornaments with 10 pixels or less marked as absent), again using rcorr.adjust().

With the ornaments we defined from our stock populations, we used functions from the R packages pedigreemm v0.3-3, sommer v4.1.0, and MCMCglmm v2.29 [48–51] to estimate heritability for size (the number of pixels), and presence/absence for individual ornaments from our pedigrees. For size, heritability was calculated using the raw data for each ornament (in the number of pixels) for all fish that had that ornament, and with fish missing that ornament marked as NA. A matrix with the relatedness between all individuals in the pedigree was calculated using the functions pedigree() and getA(). We then used the mmer() function to fit a univariate mixed effect model to estimate heritability, with a random effect - ‘animal’, which is linked to the relatedness matrix included in the model formula (ornament.size ∼ (1|animal)). For binary presence/absence data, we used a threshold of 10 pixels for presence with all sizes smaller than this marked as absent. We used the function *MCMCglmm*() to estimate heritability as implemented for binary traits [52]. We used *MCMCglmm*() for binary traits as it has been found to be more reliable for binary data than other methods available [53]. We used the priors (R = list(V = 1, fix = 1), G = list(G1 = list(V = 1, nu = 1,000, alpha.mu = 0, alpha.V = 1) with the residual variance fixed to 1 as recommended for binary data [52], and used a burnin of 100,000 and the number of iterations set to 2,500,000.

We also calculated heritability for four aggregate traits: total orange area, total black area, average orange saturation and average black saturation. Total areas were calculated respectively by summing the areas of all orange or all black ornaments. Average orange and black saturation were calculated by first finding the mean saturation of pixels in each ornament, and then calculating the weighted mean of these means (weighted by the size of each ornament). This resulted in measures for average orange and average black saturation. This method described above was used for estimating heritability of our aggregate traits.

### Strength of Y-effects on ornamentation

We tested for the potential for Y-link effects on all four aggregate traits, total orange area, total black area, average orange saturation and average black saturation, by looking at the relationship between the phenotypes of our generation three offspring and the phenotypes of their paternal and maternal grandfathers. Because the Y chromosome is passed down the patriline, grandsons will have the Y of their paternal grandfathers but not their maternal grandfathers. Therefore, we can hypothesise that if there are Y linked loci affecting total black or black area, then the slope of the paternal grandfather total black or black area should be greater than the slope of maternal grandfather phenotype on male total black or black area.

Unfortunately, we were unable to photograph and phenotype the grandfathers of our generation two offspring with our phenotype analysis pipeline. We therefore have a sample of 73 males across seven families from generation three. We carried out analyses of both maternal and paternal effects for all four aggregate traits, using mmer(). As per our heritability analysis we used a random effect - ‘animal’, which is linked to a relatedness matrix, and which controls for relatedness between individuals. We also included a fixed effect, which was either the maternal or paternal grandfather’s phenotype. The coefficients of this model represent the slope of these grandfather phenotypes (paternal and maternal) on all four aggregate traits in our generation three offspring. The formula of this analysis was: (aggregate trait ∼ Pat/Mat-GF phenotype + (1|animal)).

## Results

Following Thin Plate Spline alignment of orange and black extracted images, we identified four orange (OI - OIV) and five black ornaments (BI – BV, Fig. 1) found in our stock population. Although all orange ornaments overlapped with at least one black ornament, the overlap regions were small, and 59% of potential ornament pixels in Fig 1. were orange, 35% black, and only 6% either orange or black.

We determined correlations among ornaments in terms of size and saturation to understand the genetic relationship among different modules in our stock population. We observed generally positive correlations for both size and presence/absence among many of the orange ornaments, although many were not significant after adjusting for multiple testing (Fig. 2A, Table 1 below the diagonal, Supplementary Table 1). These correlations suggest that there may be modifier loci that globally affect the size of orange ornaments found in males, and thus also the total amount of orange pigmentation. Many previous studies that have thus used the total orange or black area as a trait may therefore have been at least in part measuring the heritability or inheritance of these loci rather than any loci related to the presence or absence of any individual ornament. In contrast, many of the correlations in size between adjacent orange and black ornaments were found to be negative (Table 1, Supplementary table 2), although few of these were significant. This perhaps indicates epistatic interactions between loci controlling ornaments, increasing the size of one ornament while reducing the size of others that are adjacent. Alternatively, it may reflect that where orange and black ornaments overlap, only the black area is visible.

**Table 1.** Strength of correlations for ornament size (below the diagonal) and saturation (above the diagonal) among orange and black ornaments. Adjusted P-value <0.05 in bold

| Ornament | OI | OII | OIII | OIV | BI | BII | BIII | BIV | BV |
| --- | --- | --- | --- | --- | --- | --- | --- | --- | --- |
| OI | - | **0.53** | 0.250 | -0.05 | -0.05 | 0.07 | -0.28 | 0.04 | -0.24 |
| OII | -0.21 | - | **0.720** | **0.40** | 0.01 | -0.03 | 0.20 | 0.21 | 0.21 |
| OIII | 0.08 | 0.19 | - | **0.41** | -0.01 | 0.03 | 0.21 | 0.17 | 0.02 |
| OIV | -0.25 | 0.28 | 0.23 | - | 0.27 | -0.10 | 0.13 | -0.01 | 0.13 |
| BI | -0.27 | -0.29 | -0.15 | 0.34 | - | 0.35 | 0.37 | 0.41 | 0.33 |
| BII | 0.13 | 0.12 | 0.01 | -0.14 | **-0.47** | - | 0.21 | **0.54** | **0.48** |
| BIII | -0.05 | 0.03 | 0.22 | 0.10 | 0.17 | -0.24 | - | 0.52 | **0.62** |
| BIV | 0.05 | **-0.44** | -0.14 | -0.07 | -0.14 | 0.02 | -0.25 | - | **0.57** |
| BV | -0.10 | -0.24 | -0.14 | 0.17 | 0.52 | -0.03 | 0.12 | 0.24 | - |

**Fig 2.**
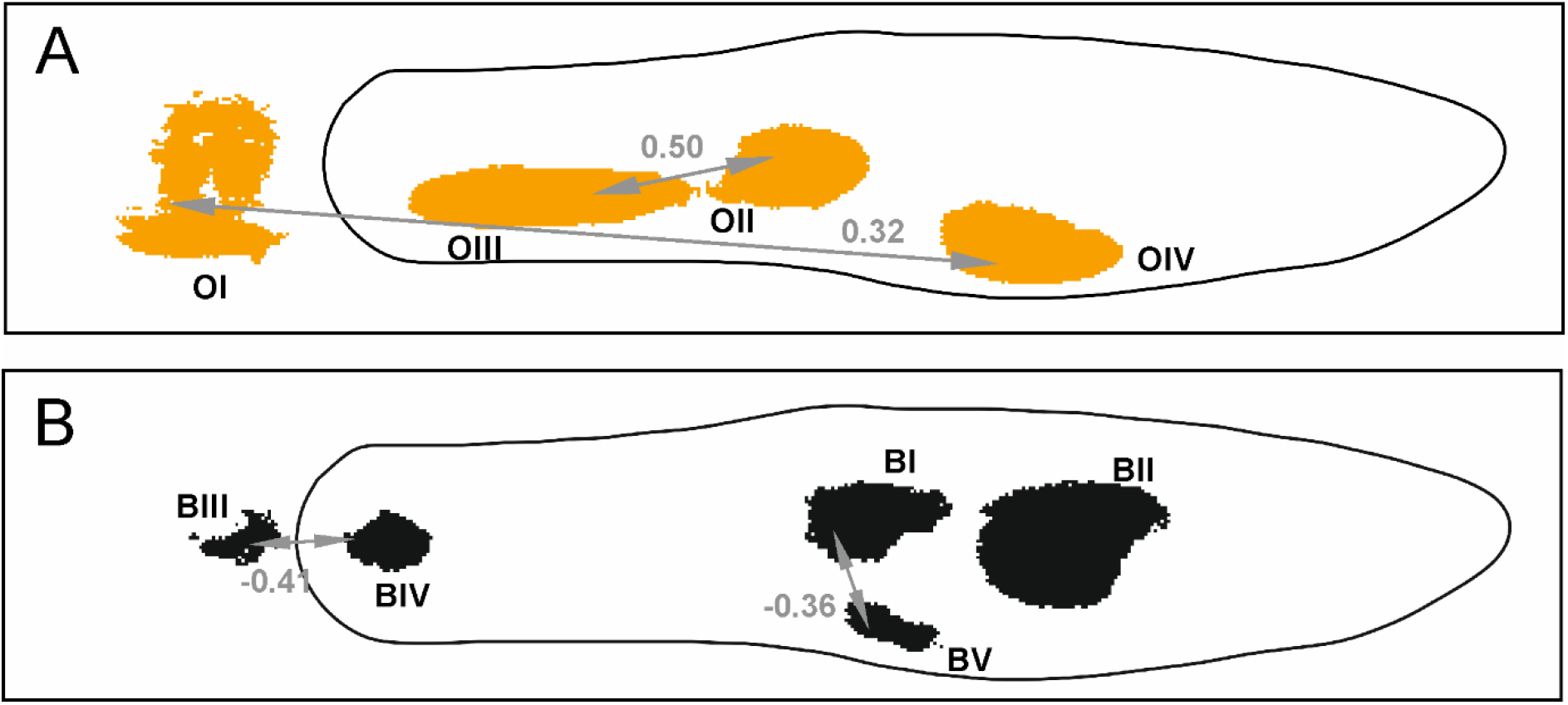
Significant correlations for presence and absence among orange (A) and black (B) ornaments. Correlations are significant to adjusted P< 0.05.

We also determined the correlation in saturation among ornaments (Table 1 above the diagonal, Supplementary Table 3). For both black and orange pigmentation, we observed generally positive correlations in saturation between adjacent ornaments of the same pigmentation type. Thus again, ornaments in the same pathway are not independent of each other, suggesting that modifier loci control saturation levels affecting multiple ornaments simultaneously. We found the correlation in saturation between orange and black ornaments was on average much lower (average = 0.049) than between ornaments of the same colour (orange average = 0.377, black average = 0.440), suggesting that modifier loci do not transcend biochemical pathways to affect both ornament types.

We used our pedigreed males to investigate the pattern of inheritance from father to son (Table 2). If the presence or absence of an ornament is exclusively Y-linked, all male offspring of a male will share their father’s phenotype. However, in cases where the presence or absence of an ornament is genetically partially or fully under autosomal, X or pseudoautsomal control, sons will frequently have different phenotypes than their fathers.

**Table 2.**
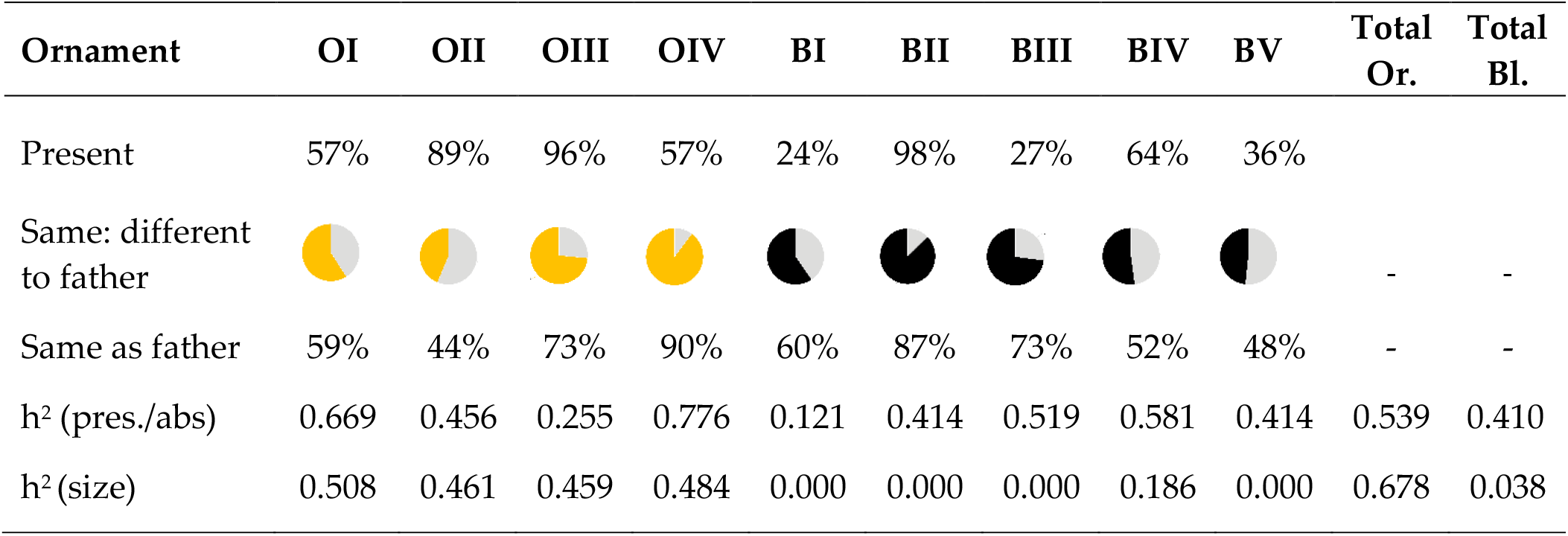
Father-son heritability of presence/absence and area for each ornament, and for total orange and total black area.

| Ornament | OI | OII | OIII | OIV | BI | BII | BIII | BIV | BV | Total Or. | Total Bl. |
| --- | --- | --- | --- | --- | --- | --- | --- | --- | --- | --- | --- |
| Present | 57% | 89% | 96% | 57% | 24% | 98% | 27% | 64% | 36% |  |  |
| Same: different to father |  |  |  |  |  |  |  |  |  | - | - |
| Same as father | 59% | 44% | 73% | 90% | 60% | 87% | 73% | 52% | 48% | - | - |
| $h^2$ (pres./abs) | 0.669 | 0.456 | 0.255 | 0.776 | 0.121 | 0.414 | 0.519 | 0.581 | 0.414 | 0.539 | 0.410 |
| $h^2$ (size) | 0.508 | 0.461 | 0.459 | 0.484 | 0.000 | 0.000 | 0.000 | 0.186 | 0.000 | 0.678 | 0.038 |

Our results show that no ornament displays the inheritance pattern expected for a purely male specific Y-linked trait. For two ornaments (OII and BV), we observe a majority of male offspring have the opposite phenotype to their father. Overall these results suggest that individual orange and black ornaments are not linked to the Y chromosome in our population.

Most ornaments show high heritability except for ornaments OIII and BI (Table 2). In contrast, heritability for total area differed markedly between orange (0.678) and black (0.038) ornaments. We also found heritable genetic variation for average (across ornaments) orange saturation (h2 = 0.232) and for average black saturation (h2 = 0.607).

Our analysis of paternal and maternal grandfather effects (Table 3) identified a stronger relationship between males and their paternal grandfather compared to their maternal grandfather for total orange area and orange saturation. This indicates that there may indeed be Y-linked modifier loci that affect total orange area and average orange saturation in guppies, and is consistent with our previous analyses which show positive correlations in size and saturation among orange ornaments. Although there was a greater coefficient for total black area between males and their paternal compared to maternal grandfather, the coefficients for black saturation were negative for both grandfathers.

**Table 3.** Coefficients from our analysis of paternal and maternal grandfather effects on aggregate offspring phenotypes.

|  | Paternal GF Effect | Maternal GF Effect |
| --- | --- | --- |
| Total Orange Area | 0.807 | 0.232 |
| Total Black Area | 0.452 | -0.214 |
| Average Orange Saturation | 0.313 | -0.186 |
| Average Black Saturation | -0.465 | -0.056 |

## Discussion

Our study uses high resolution image analysis to define individual ornaments while reducing human bias, in order to examine inheritance patterns across a dataset of 214 offspring from 19 families. By computationally clustering pixels automatically in order to define individual ornaments [36], we provide a comprehensive and agnostic diagnostic method for colouration patterning in a semi-natural outbred population. Using this method, we identified four orange and five black ornaments (Fig. 1) in our population that we then used to study modulation, heritability and mode of inheritance.

### Modulation of ornamentation

The colourful ornaments found in animals are often treated as independent phenotypes, with the relationships among different ornaments less well considered [54]. However, in many cases the functions of these ornaments are not independent, but as whole patterns in conjunction [55]. Furthermore, some ornaments, particularly those that share the same biochemical basis, may have shared aspects of genetic architecture through the pathways that control them, and so treating them as individual units may not always be appropriate.

The colour patterns of guppies are used as a sexual signal. It has been proposed that such signals might be made more conspicuous, and therefore attractive to females, by greater differences between adjacent patches of colour [54]. If this is the case for guppies, we might expect selection for linkage between adjacent black and orange patches. We tested this by examining correlations in presence and absence between different male ornaments (Table 1). Although we found some strong positive correlations for size within ornaments of the same colour, there was little correlation among adjacent ornaments of different colours (Supplementary Table 2). Our results therefore do not provide evidence of linkage between adjacent and contrasting male ornaments in our population of guppies.

Although we find little correlation between ornaments from different biochemical pathways, we do find several lines of evidence that guppy ornaments of the same colour are not independent of each other. Interestingly, orange and black ornaments displayed different genetic characteristics, with high heritability for size (Table 2) and high correlations in size (Table 1, Fig. 2) between many of the orange ornaments, but not black ornaments. However, for both black and orange pigmentation, we observed generally positive correlations in saturation between adjacent ornaments of the same pigmentation type (Table 1). This suggests the presence of one or more global modifier loci affecting saturation levels and total size for multiple ornaments simultaneously, while other independent loci encode for the presence or absence of individual ornaments. Furthermore, the higher similarity between grandsons and their paternal grandfathers compared to maternal grandfathers in ornament size and orange saturation (Table 3) suggests a Y-linked component to this modifier locus, discussed below.

### Sex linkage

Many guppy colour traits have been shown to be Y-linked, reviewed by Lindholm and Breden [56] and Kottler and Schartl [57]. However, much of this work has either been done on ornaments in domestic guppy breeds and therefore unlikely to be found in the wild [31,32], or on composite traits, such as aggregate colour area [21,33,34]. We assessed the potential for Y-linkage in individual ornaments from our semi-natural populations under the expectation that any Y-linked ornament displayed by a father would be also present in all his sons. Despite this previous work, we did not observe Y inheritance patterns for presence or absence of any single one of our nine ornaments. This suggests that individual male colour patterns in guppies from our population are not Y-linked. This is also reflected in the heritability for presence/absence of these ornaments, with more than half of ornaments with heritability <0.5. Even for the ornaments inherited with the highest heritability (OIV), 9.7% of offspring males display a different phenotype to their father.

Recent studies have suggested that the guppy Y chromosome is divided into a relatively small region where recombination between the X and Y never occurs [58–61], and a larger adjacent region where recombination is rare, occurring in between 1% and 0.1% of males [59,62,63]. However, because the vast majority of recombination in males is confined to the telomeric ends of the sex chromosome, nearly the entire chromosome is passed on as a Y chromosome in >99% of male offspring. Therefore, rare X-Y recombination events cannot explain the difference between paternal and filial phenotypes we observe in our ornaments.

These results are at first perplexing in light of previous work, and there are several potential explanations. First, it is possible that our population has lost Y-linkage of ornaments in the >20 years since it was collected from the wild. However, we think this is not terribly likely given the large size of the founding colony and the fact that it has been subsequently maintained as an outbred, freely mating population. Both of these strategies have maximized genetic diversity and minimized potential bottlenecks. Moreover, Y-linkage has been observed in other lab and inbred populations [21,64], and it is not clear how or why Y-linkage would erode.

Second, considerable variation in Y-linkage exists among wild Trinidadian populations [65], with notable differences between high predation and low predation populations in particular [33,34,59,66]. It is therefore possible that difference between our results and others may simply be due to differences between populations. Although we also think this is unlikely given that the Quare population upon which our stock population is founded has been studied using aggregate methods, described below [34], it is however is important for future studies to quantify Y-linkage of individual ornaments in multiple populations.

However, our results are consistent with studies which have characterized aggregate ornamental area, which often identify significant Y contributions. For example, Postma *et al* [35] and Brooks and Endler [21] both identified significant levels of Y-linked genetic variation when assessing total ornamental areas. Our analysis suggests that although the presence or absence of individual ornaments are not often Y-linked in natural populations, the Y chromosome does carry one or more modifier loci that affect many ornaments in aggregate (Table 3). Therefore when studies use the total orange or black area as a trait they are in large part measuring the mode of inheritance of these loci rather than any loci related to the presence or absence of any individual ornament [21,34].

It may be that these Y-linked modifier loci work through a simple mechanism, such as testosterone level or testosterone sensitivity, which has been shown to affect total ornament area [32–34]. Indeed, the extensive variation exhibited in total ornament area among natural [34] and introduced [33] populations hints at a simple genetic architecture, as the alternative of rapid and repeated movement of many underlying genes to the relatively small Y chromosome, is somewhat improbable. Given the importance of testosterone level and sensitivity in fish growth rates, maturation and colouration [67–69], and the variation observed in natural guppy populations in these traits [70], it is likely that testosterone and/or testosterone sensitivity vary across populations, and this may interact with colouration. The nature of these interactions, and the identification of the loci responsible for these Y-linked effects remain interesting areas of future research.

### Conclusion

Surprisingly, we did not observe inheritance patterns for presence and absence for any of our nine ornaments that were consistent with Y-linkage. However, we did find evidence of Y-linked modifier loci that affect the aggregate phenotypes of total orange and black area. Given the results of our study, the Y-linkage of individual male colour traits may therefore not be as important in the rapid evolution of colour changes in natural populations of guppies [33] or in the expansions of the non-recombining Y region [30] as previously thought. However, Y-linked modifier loci that affect aggregate traits may well play a role. Alternatively other mechanisms to resolve sexual conflict, such as changes in gene regulation [71] may alone be enough to explain colour dimorphism in this species.

## Acknowledgements

This work was supported by a grant from the European Research Council (grant agreement N0. 680951) to JEM, who also gratefully acknowledges support from a Canada 150 Research Chair and NSERC. We thank Natasha Bloch for her early input, and Ben Furman, Jacelyn Shu, David Metzger, Ben Sandkam, Yuying Lin, Lydia Fong, Pedro Almeida and Alberto Corral-Lopez for their helpful comments.

