## Supplementary figures and tables for "HIGH-RESOLUTION CHARACTERIZATION OF MALE ORNAMENTATION AND REEVALUATION OF SEX LINKAGE IN GUPPIES"

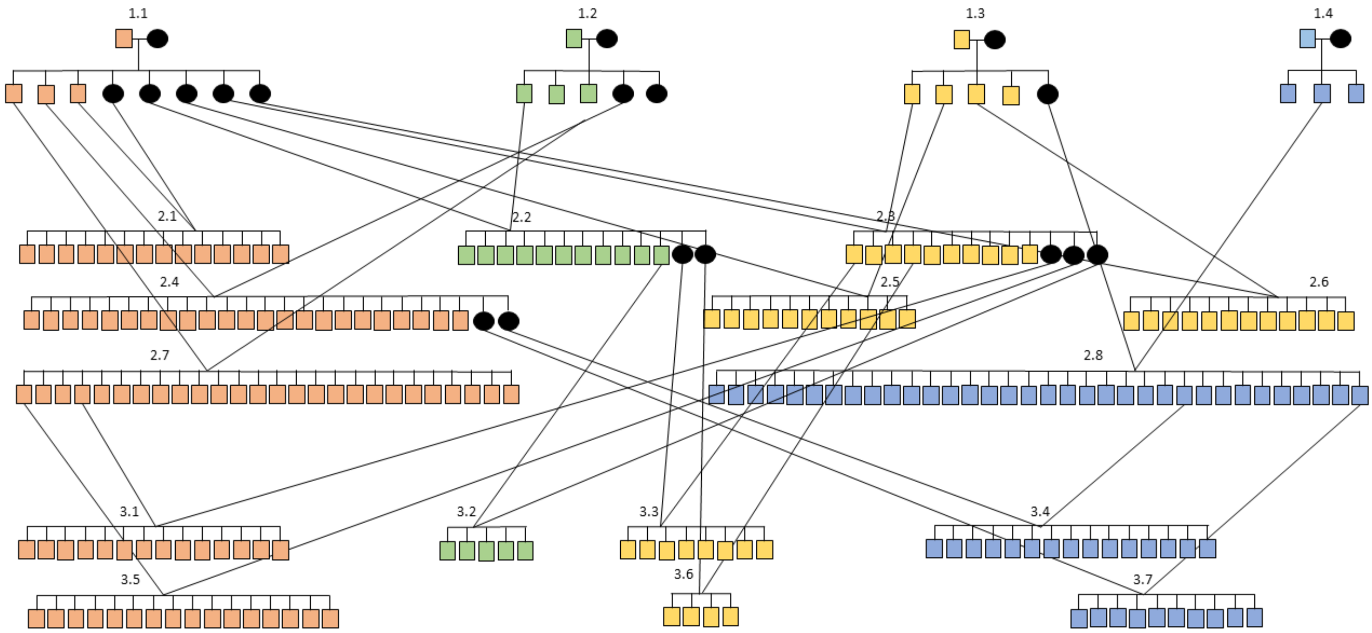


**Supplementary Fig. 1. Crossing design for pedigrees with generations 1-3. All phenotyped males and females used for breeding are shown.**

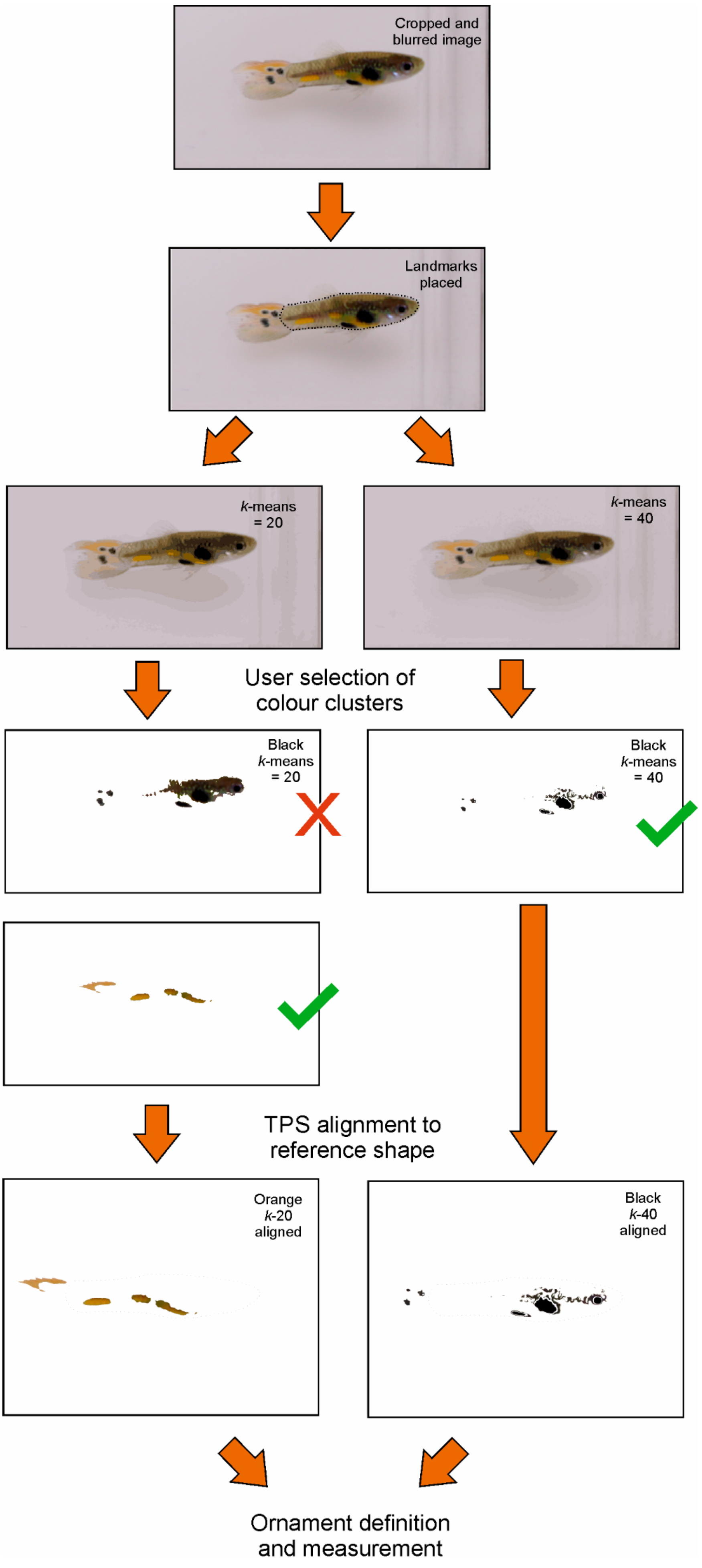
Cropped and blurred image
Landmarks placed
k-means = 20
k-means = 40
User selection of colour clusters
Black k-means = 20
Black k-means = 40
TPS alignment to reference shape
Orange k-20 aligned
Black k-40 aligned
Ornament definition and measurement

**Supplementary Fig. 2. Overview of image analysis pipeline steps and outputs.**

**Supplementary Table 1. Strength of correlations for ornament presence/absence, among carotenoid and melanic ornaments**. Below diagonal; adjusted P-value (significant values P <0.05 in bold), and above diagonal; the correlation coefficient.

| Ornament | OI | OII | OIII | OIV | BI | BII | BIII | BIV | BV |
|---|---|---|---|---|---|---|---|---|---|
| OI | | -0.03 | 0.14 | **0.32** | -0.1 | -0.02 | 0.02 | -0.04 | 0 |
| OII | 1.0000 | | **0.5** | 0.07 | 0.03 | -0.1 | 0.16 | -0.02 | -0.08 |
| OIII | 1.0000 | **<.0001** | | 0.1 | -0.11 | -0.11 | -0.05 | 0.09 | 0.02 |
| OIV | **0.0005** | 1.0000 | 1.0000 | | 0.05 | 0.03 | **0.36** | -0.08 | 0.05 |
| BI | 1.0000 | 1.0000 | 1.0000 | 1.0000 | | -0.07 | 0.2 | -0.19 | **-0.36** |
| BII | 1.0000 | 1.0000 | 1.0000 | 1.0000 | 1.0000 | | -0.02 | 0.17 | 0.01 |
| BIII | 1.0000 | 0.9725 | 1.0000 | **<.0001** | 0.3046 | 1.0000 | | **-0.41** | -0.04 |
| BIV | 1.0000 | 1.0000 | 1.0000 | 1.0000 | 0.3701 | 0.7538 | **<.0001** | | 0.05 |
| BV | 1.0000 | 1.0000 | 1.0000 | 1.0000 | **<.0001** | 1.0000 | 1.0000 | 1.0000 | |

**Supplementary Table 2. Strength of correlations for ornament size, among carotenoid and melanic ornaments.** Below diagonal; adjusted P-value (significant values P <0.05 in bold), and above diagonal; the correlation coefficient.

| Ornament | OI | OII | OIII | OIV | BI | BII | BIII | BIV | BV |
|---|---|---|---|---|---|---|---|---|---|
| OI | - | -0.21 | 0.08 | -0.25 | -0.27 | 0.13 | -0.05 | 0.05 | -0.1 |
| OII | 1.0000 | - | 0.19 | 0.28 | -0.29 | 0.12 | -0.03 | **-0.44** | -0.24 |
| OIII | 1.0000 | 0.8722 | - | 0.23 | -0.15 | 0.01 | -0.22 | -0.14 | -0.14 |
| OIV | 1.0000 | 0.5714 | 1.0000 | - | 0.34 | -0.14 | 0.1 | -0.07 | 0.17 |
| BI | 1.0000 | 0.6371 | 1.0000 | 0.7366 | - | **-0.47** | 0.17 | -0.14 | 0.52 |
| BII | 1.0000 | 1.0000 | 1.0000 | 1.0000 | **0.0039** | - | -0.24 | 0.02 | -0.03 |
| BIII | 1.0000 | 1.0000 | 1.0000 | 1.0000 | 1.0000 | 1.0000 | - | 0.16 | 0.12 |
| BIV | 1.0000 | **0.0254** | 1.0000 | 1.0000 | 1.0000 | 1.0000 | 1.0000 | - | 0.24 |
| BV | 1.0000 | 1.0000 | 1.0000 | 1.0000 | 1.0000 | 1.0000 | 1.0000 | 1.0000 | - |

**Supplementary Table 3. Strength of correlations for ornament saturation, among carotenoid and melanic ornaments**. Below diagonal; adjusted P-value (significant values P <0.05 in bold), and above diagonal; the correlation coefficient.

| Ornament | OI | OII | OIII | OIV | BI | BII | BIII | BIV | BV |
|---|---|---|---|---|---|---|---|---|---|
| OI | - | **0.53** | 0.25 | -0.05 | -0.05 | 0.07 | -0.28 | 0.04 | -0.24 |
| OII | **0.0004** | - | **0.72** | **0.4** | 0.01 | -0.03 | 0.2 | 0.21 | 0.21 |
| OIII | 0.6668 | **<.0001** | - | **0.41** | -0.01 | 0.03 | 0.21 | 0.17 | 0.02 |
| OIV | 1.0000 | **0.0085** | **0.0019** | - | 0.27 | -0.1 | 0.13 | -0.01 | -0.13 |
| BI | 1.0000 | 1.0000 | 1.0000 | 1.0000 | - | 0.35 | 0.37 | 0.41 | 0.33 |
| BII | 1.0000 | 1.0000 | 1.0000 | 1.0000 | 0.0548 | - | 0.21 | **0.54** | **0.48** |
| BIII | 1.0000 | 1.0000 | 1.0000 | 1.0000 | 0.2563 | 1.0000 | - | 0.52 | **0.62** |
| BIV | 1.0000 | 1.0000 | 1.0000 | 1.0000 | 0.5327 | **<.0001** | 0.6668 | - | **0.57** |
| BV | 1.0000 | 1.0000 | 1.0000 | 1.0000 | 1.0000 | **0.0031** | **0.0056** | **0.0141** | - |
